# Early-life sleep disruption impairs subtle social behaviors during pair bond formation in prairie voles: A computer vision study

**DOI:** 10.1101/2022.10.12.511931

**Authors:** Lezio S. Bueno-Junior, Carolyn E. Jones-Tinsley, Peyton T. Wickham, Brendon O. Watson, Miranda M. Lim

## Abstract

Early life sleep disruption (ELSD) has been shown to have long lasting effects on social behavior in adult prairie voles (*Microtus ochrogaster*), including impaired expression of pair bond behavior during partner preference testing (e.g., reduced huddling with a pair bonded partner). However, due to the limitations of manual behavior tracking, the behavioral effects of ELSD across the entire time course of pair bond formation have not yet been described, hindering our ability to trace mechanisms. Here, we used computer vision to precisely track multiple behaviors during opposite-sex cohabitation of prairie voles. Male-female pairs were allowed to interact through a mesh divider in the home cage for 72 h, providing variables of body direction, distance-to-divider, and locomotion speed, with temporal resolution as high as the video frame rate (20-25 Hz). We found that control males displayed periodic, stereotyped patterns of body orientation towards females during pair bond formation. In contrast, ELSD males showed reduced duration and ultradian periodicity of these body orientation behaviors towards females. Furthermore, in both sexes, ELSD altered stereotypical spatial and temporal patterns of locomotion seen in control animals that typically varied across days of cohabitation and light/dark periods. This study highlights the utility of computer vision in deep characterization of subtle behaviors and allows a more comprehensive behavioral assessment of the profound and persistent effects of ELSD on later life social behavior. Our findings may shed light on causal mechanisms underlying human neurodevelopmental disorders featuring sleep disruption and social deficits, such as autism spectrum disorder.

## INTRODUCTION

Early life sleep changes are prominent in disorders such as autism spectrum disorders (ASD) that feature prominent social alterations later in life^1–4^. Prairie voles (*Microtus ochrogaster*) are a socially monogamous wild rodent species that are an ideal animal model in which to study the relationship of early life sleep to adult social behavior^5–7^. Unlike traditional lab rats and mice, prairie voles show strong affiliations with opposite-sex mates after a period of cohabitation. This so-called “partner preference” behavior is used as a proxy for pair bonding described in the wild^8–11^. Furthermore, unlike lab rats and mice, prairie voles display side-by-side huddling with conspecifics for prolonged periods of time, often choosing huddling over exploration in a novel environment^12^. Thus, voles are uniquely suited to dissect the nuanced and complex locomotor aspects of social bonding, many of which may be relevant to aspects of human neurodevelopmental disorders, including ASD.

Normally, prairie voles require between 6 and 72 hours of cohabitation with an opposite sex mate in order to form a pair bond – the duration of which may vary depending on several reported factors, including sex, various stressors, hormonal status, and presence of mating^9,13–15^. Recently, we have shown that early life sleep disruption (ELSD) directly alters social behavior between opposite-sex mates in adulthood. Specifically, ELSD during the 3^rd^ postnatal week in prairie vole pups causes adult males (∼100 days old) to reduce huddling behavior with their female partner during the partner preference test – the proxy for assessing the strength of a pair bond^16^. Intriguingly, social memory and selective aggression towards the stranger are preserved, indicating that effects of ELSD seem to be preferentially biased towards affiliative interactions^16^. However, our understanding of effects of ELSD on affiliative behavior have been limited due to prior methods of utilizing manual behavior tracking.

Herein, we report an innovative and unbiased approach to characterize locomotor behavior during pair bond formation in prairie voles using computer vision and customized analyses. Using a 72-hour cohabitation protocol with a male and female prairie vole pair separated by a mesh divider, we perform deep phenotyping to identify new, subtle changes in social behavior during the entirety of the time span of pair bond formation in both sexes. Two strategies are proposed in combination to examine the behavior of prairie vole pairs while they interact through mesh dividers. One strategy utilizes the spatial dimension via overhead video, so that male and female individuals can be tracked in their respective 2D spaces without spatial overlap, generating well-controlled measures of within-pair interactions. The second strategy utilizes the temporal dimension across 72 hours with enough temporal resolution to capture social dynamics, such as the latency with which the animals habituate to each other, as well as the incidence of social interactions over time.

This study provides a description of a method that examines both temporal and spatial dimensions of pair bond formation without requiring *a priori* temporal or spatial discretization. Our results corroborate prior reported findings that ELSD males show more profoundly impaired affiliative behavior^16^, but we also discovered new, subtle social phenotypes across both sexes with regard to both the timing and spatial topography of within-pair interactions.

## RESULTS

### Early life sleep disruption and video recording later in adulthood

ELSD phenotypes from both sexes were produced by exposing prairie vole litters to gentle orbital shaking of the home cage from postnatal day (P) 14 to P21 (Figure 1A), in accordance with our previous study^16^. This method has been shown to reduce REM sleep and fragment NREM sleep in prairie voles without affecting parental care or stress hormones. Same-sex siblings were then co-housed without experimental manipulations until adulthood (∼P100).

**Figure 1.**
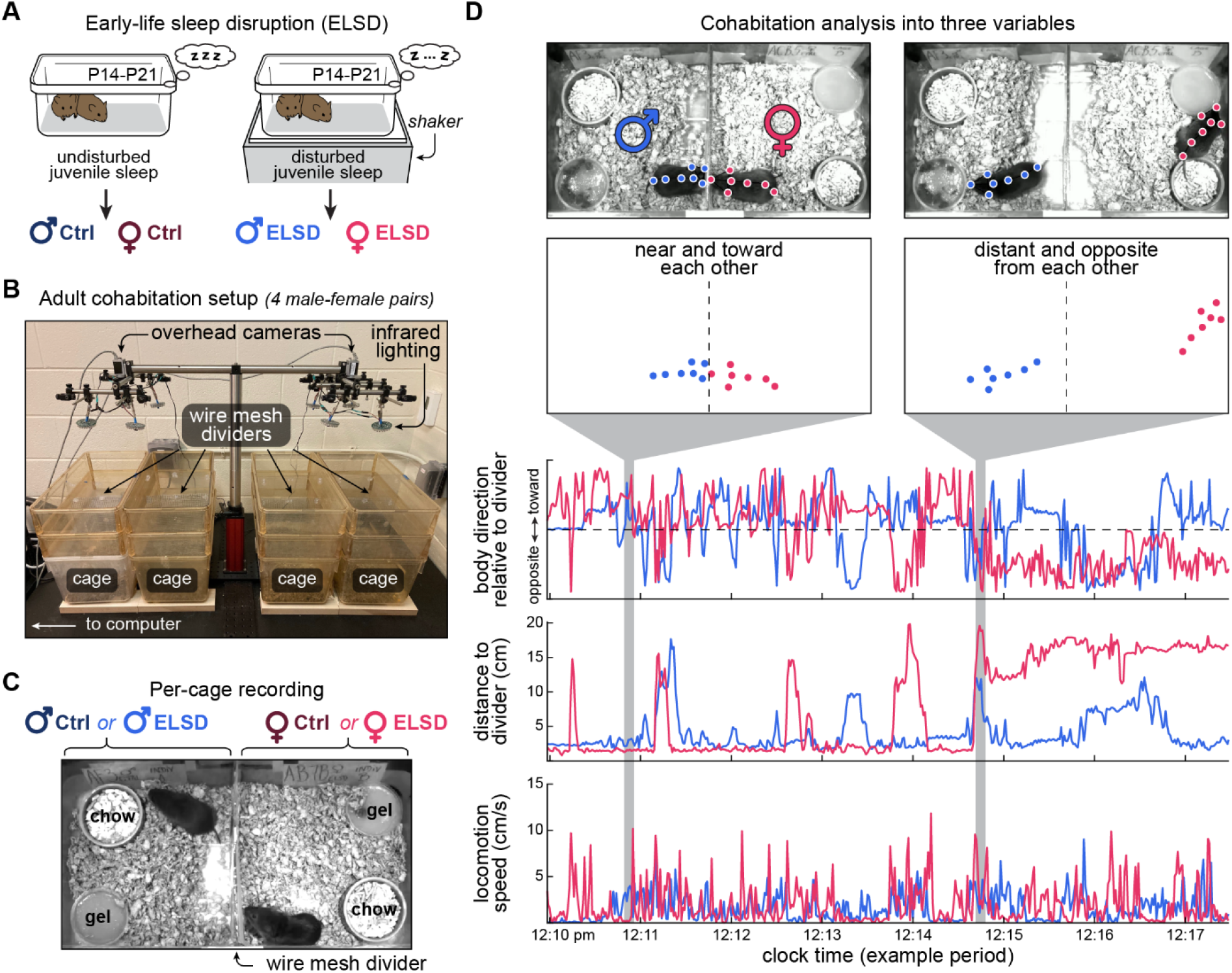
Juvenile sleep disruption and adult video recording. (A) Prairie vole litters were exposed to gentle cage agitation during a sensitive developmental window (P14-P21), resulting in early-life sleep disruption, or ELSD^16^. (B-C) Once adults, ELSD animals (or their controls) were assigned to mutually naive male-female pairs. These pairs were monitored from an overhead angle for 72 h under circadian light/dark switching. (D) Using DeepLabCut^17^ on video from each animal individually, animals were tracked as if they were moving arrows, before being re-combined into pairs. Behavioral tracking data were then processed into standardized measures across recordings: body direction, distance from divider and locomotion speed.

After reaching adulthood, prairie voles were filmed using an overhead video setup with two infrared-sensitive cameras (Figure 1B). It was designed to record four male-female pairs simultaneously over 72 h in a room with circadian light/dark switching. Infrared illumination with infrared-only camera filters kept video brightness constant despite the light/dark switching. Male-female pairs were assigned randomly into all possible ELSD and control (Ctrl) combinations: Male-ELSD/Female-ELSD, Male-ELSD/Female-Ctrl, Male-Ctrl/Female-ELSD, Male-Ctrl/Female-Ctrl (Figure 1C). Individuals were naive to each other at the beginning of recordings and were allowed to interact through lab-made wire mesh dividers. Chow and gel were placed consistently across recordings, always relative to the animal (Figure 1C).

Using DeepLabCut (DLC)^17^, we tracked each animal using markers on the nose, ears and centerline of the body (Figure 1D, top). We then converted DLC data from pixels to centimeters and obtained three behavioral measurements per video frame: body direction relative to divider, distance to divider and locomotion speed (Figure 1D, bottom; see Methods). These measures revealed novel details on within-pair interactions, suggesting a spectrum with mutual social exploration on one extreme and mutual indifference on the other (Figure 1D; Movie S1). We quantified this as described next.

### Male-ELSD prairie voles exhibit lower orientation towards females in the initial 12 h of cohabitation

Figure 2A shows behavioral measures in 1 h bins, across the 72-h cohabitation. First, we compared males vs. females irrespective of ELSD or Ctrl treatments. Across time bins, we observed that males were constantly more likely to behave toward the divider (Figure 2A, left graph) (effect of sex grouping: F_1,3816_ = 3.0*10^3^, P = 0; group vs. time interaction: F_71,3816_ = 1.431, P = 0.011) and females showed higher locomotion speed, particularly in the initial hours of recording (Figure 2A, right graph) (effect of sex grouping: F_1,3816_ = 169.100, P = 7.3*10^−38^). This suggests that males spent more time investigating the partner, whereas females were more likely to roam across the home cage area, especially earlier during cohabitation. Also, across sexes, we observed a gradual increase in distance to divider (Figure 2A, center graph) (effect of time: F_71,3816_ = 3.257, P = 3.9*10^−18^) and gradual decrease in locomotion speed (Figure 2A, right graph) (effect of time: F_71, 3816_ = 10.994, P = 2.2*10^−107^). This suggests a transition from higher activity near the divider on Day 1 (possibly due to novelty) to lower activity in a broader home cage area in later periods (possibly due to habituation).

**Figure 2.**
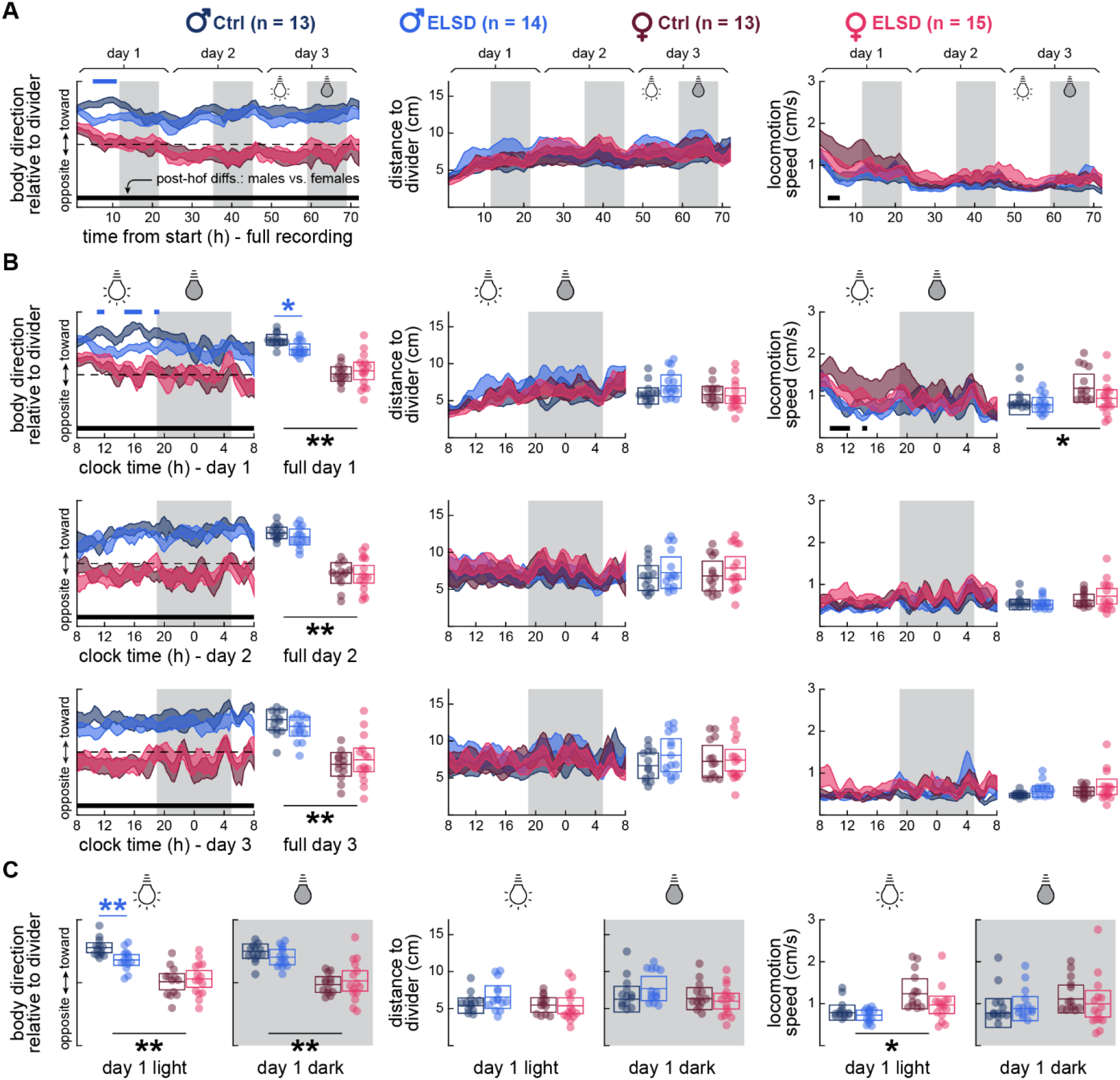
Male-ELSD prairie voles exhibit lower orientation towards females in the initial 12 h of cohabitation. (A) Behavioral measures across 72 h cohabitation in 1 h bins. Light/dark periods are indicated with background shading. Shaded curves represent standard errors around means. Both male groups were more likely than females to orient toward the divider throughout the experiment (left graph) and both female groups showed higher locomotor activity than males in the initial hours of recording (right graph), as indicated by post-hoc differences (black bars on bottom). All groups showed a tendency from high activity near the divider on Day 1 to lower activity in a broader home cage area in later periods. Male-ELSD animals were less likely than Male-Ctrl to orient toward the divider during the initial 12 h of recording (blue bar on top). Post-hoc differences: P < 0.01. (B) Curve graphs: same data, but in 20 min bins and divided into days of recording. Male-ELSD animals were indeed less likely than Male-Ctrl to behave toward the divider. Box plots: averages from the corresponding curve graphs, reinforcing the difference between male groups (blue asterisks), as well as between-sex differences (black asterisks). (C) Averaged box plot data like (B) but focused on Day 1. The Male-ELSD effect of (B) coincided with the light period of Day 1. *P < 0.05. **P < 0.005.

Then, we examined ELSD vs. Ctrl animals per sex. During the initial 12 h of recording, Male-ELSD animals were less likely to orient toward the divider than Male-Ctrl (Figure 2A, left graph) (effect of ELSD vs. Ctrl grouping: F_1,1800_ = 67.918, P = 3.2*10^−16^). To further specify this difference, we performed ELSD vs. Ctrl comparisons at higher temporal resolution (20 min bins) and in separate 24-h segments aligned with clock time, as shown in Figure 2B. When analyzing Day 1 alone, we found stronger body direction differences between Male-ELSD and Male-Ctrl (effect of ELSD vs. Ctrl grouping: F_1,1800_ = 126.226, P = 2.3*10^−28^; group vs. time interaction: F_71,1800_ = 1.345, P = 0.031). We also averaged across the time bins of Day 1 per animal and again found a significant body direction difference between the two male groups (see box plots; F_1,25_ = 5.782, P = 0.024). Differences between sexes irrespective of ELSD or Ctrl treatments were again present in body direction (effect of sex grouping: F_1,3816_ = 1.5*10^3^, P = 4.2*10^−283^; box plots: F_1,53_ = 58.634, P = 4.0*10^−10^) and locomotion speed (effect of sex grouping: F_1,3816_ = 180.303, P = 3.4*10^−40^; box plots: F_1,53_ = 4.810, P = 0.033). This confirms the sex-specific behaviors we outlined above, i.e., slower activity toward the partner in males, faster activity across the home cage in females during Day 1. Figure 2C focuses further on Day 1 by separately averaging its light and dark periods. During the light phase, there was an even stronger difference between the male groups (F_1,25_ = 10.961, P = 0.003). This difference coincides with the period of novelty-induced home-cage activity explained above (Figure 2A), suggesting a relationship between environmental novelty and altered Male-ELSD body direction (circadian influences not necessarily involved; see Discussion).

Therefore, Figure 2 demonstrates an optimal time interval and behavioral measure to capture ELSD effects on male prairie voles, in addition to demonstrating sex differences in body direction and locomotion speed irrespective of ELSD treatment. These findings suggest that ELSD preferentially impairs the social components of male behavior in the initial hours of cohabitation.

### ELSD affects body direction rhythms in sex-specific manners

Interestingly, the curve plots in Figure 2B also show somewhat regular fluctuations across all behavioral measures and groups of animals. These fluctuations suggest cycles in the timescale of around 2-3 h, i.e., ultradian rhythms, especially during dark periods. We address this in Figure 3.

**Figure 3.**
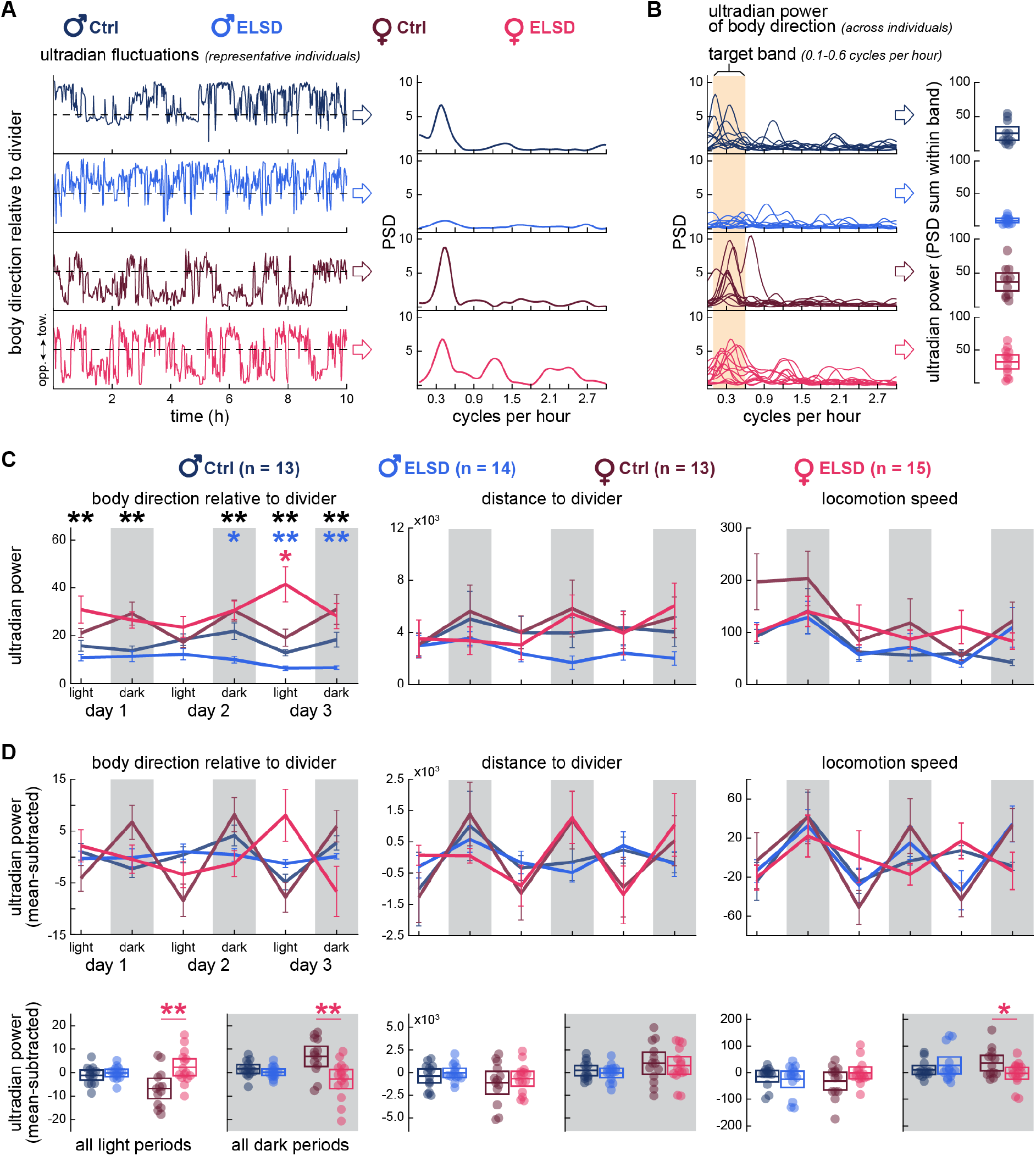
ELSD affects body direction rhythms in sex-specific manners. (A) Left: 10-h curves (1-s bins) from representative individuals per group, showing body direction fluctuations recurring every 2-3 hours. Right: power spectral density (PSD) of such fluctuations, showing peaks at around 0.4 cycles per hour. The PSD peak was lower in the Male-ELSD individual. (B) Left curves: PSD curves for all animals in each group, showing that PSD peaks were indeed lower in Male-ELSD animals as a group. By summing PSD values within the frequency band containing most of PSD peaks (0.1-0.6 cycles/h) we obtained band power values per individual, i.e., “ultradian power” of body direction. Right box plots: ultradian power of body direction across groups, specifically during the dark period of Day 2. (C) Same analysis as in (B), but across behavioral measures and recording periods. Females showed generally stronger ultradian periodicity of body direction than males (black asterisks). Male-ELSD animals showed weaker ultradian periodicity in body direction than Male-Ctrl from Day 2 (blue asterisks). Female-Ctrl animals seemed to show a circadian fluctuation in the ultradian power of body direction, and this effect was not as apparent in Female-ELSD. We analyzed this more deeply in (D). (D) Circadian fluctuations of ultradian power were magnified by mean-subtracting the data in (C). Light and dark periods were then separately averaged and depicted as box plots. This revealed a female-specific effect of ELSD: a disruption of both ultradian and circadian behavioral rhythms, especially in the body direction variable. *P < 0.05. **P < 0.005.

Figure 3A shows fluctuations in body direction over a 10-h period for one representative individual per group of animals. It also shows the power spectral density (PSD) of those fluctuations, and which frequencies were most prevalent. PSD peaked at around 0.4 cycles per hour, with strong peaks in all individuals except the Male-ELSD one. Figure 3B exhibits PSD curves from all individuals per group, illustrating that PSD peaks appeared indeed weaker in Male-ELSD animals. We quantified this by summing the 0.1-0.6 cycle/h PSD values per individual, as exemplified in the box plots of Figure 3B. Here we will refer to 0.1-0.6 cycle/h as the “ultradian band”, corresponding to cycles of behavioral activity recurring every 2-3 h approximately.

The ultradian power analysis exemplified in Figure 3B is from body direction during the dark period of Day 2. The same analysis is shown in Figure 3C, but across behavioral variables and time periods. Particularly in the body direction variable, females were more likely than males to exhibit ultradian periodicity throughout the 72-h cohabitation, irrespective of ELSD or Ctrl status (Figure 3C, black asterisks) (effect of sex grouping: F_1,318_ = 84.807, P = 4.6*10^−18^). This indicates that females are more likely than males to reorient themselves every 2-3 h. Within-sex effects of ELSD on body direction were confined to later cohabitation periods: Male-ELSD showed constantly weaker ultradian power, especially starting on Day 2 (effect of ELSD vs. Ctrl grouping: F_1,150_ = 35.740, P = 1.6*10^−8^). This indicates that ELSD further reduces the tendency of males to reorient themselves every 2-3 h, especially as they habituate to the environment.

Figure 3C also shows a female-specific effect of ELSD on body direction. First, we noticed that Female-Ctrl animals show a fluctuation in ultradian power across periods: weaker in light, stronger in dark. We then noticed a disruption of such pattern in Female-ELSD, including a localized increase in ultradian power during the light period of Day 3 (effect of ELSD vs. Ctrl grouping: F_1,156_ = 3.925, P = 0.049). This suggested an ELSD-sensitive relationship between the ultradian and circadian timescales, which we investigated more deeply in Figure 3D. Ultradian power curves of each animal were subtracted by their own moving averages. This resulted in the detrended curves of Figure 3D, with clearer light/dark cycling. We then averaged light and dark periods separately per animal, producing the box plots of Figure 3D. The strongest effects were found in the ultradian power of body direction. More specifically, Female-Ctrl animals were observed to alternate between lower and higher ultradian power during light and dark periods, respectively. This pattern was disrupted in Female-ELSD (Figure 3D, bottom left) (light period: F_1,26_ = 10.265, P = 0.004; dark period: F_1,26_ = 10.588, P = 0.003). Female-ELSD additionally showed a slightly lower ultradian power of locomotion speed during dark periods than controls (Figure 3D, bottom right) (F_1,26_ = 4.327, P = 0.047).

We found no within-sex ELSD effects on the ultradian periodicity of distance to divider or locomotion speed across periods (Figure 3C). This was also true for light vs. dark comparisons, except for the relatively weak effect in female locomotion speed described above (Figure 3D). This suggests that ELSD preferentially impairs body direction rhythms in sex-specific manners, while sparing more general aspects of home cage roaming, consistent with Figure 2.

### ELSD alters both spatial occupancy and body orientation in sex-specific manners

We next mapped body direction data onto the cohabitation cage area, resulting in the spatial patterns of Figure 4 (2 mm grids). The heatmaps in Figure 4A were produced by Z scoring across male animals and recording periods, and then separately averaging across Male-Ctrl or Male-ELSD animals per recording period. During the light period of Day 1, we observed an area near and along the divider where animals spent more time orienting toward the divider, especially Male-Ctrl animals (Figure 4A; see red areas within 5 cm from the divider). To quantify this, the heatmap of each individual animal was averaged vertically into a body direction vs. distance to divider curve. Standard errors from such curves are presented as the bottom graphs of Figure 4A. During the light period of Day 1, we observed that Male-Ctrl animals indeed spent more time near and toward the divider (effect of ELSD vs. Ctrl grouping: F_1,2500_ = 43.919, P = 4.2*10^−11^; group vs. distance interaction: F_99,2500_ = 2.816, P = 1.1*10^−17^). With this, we clarify the spatial pattern behind the body direction results of Figure 2.

**Figure 4.**
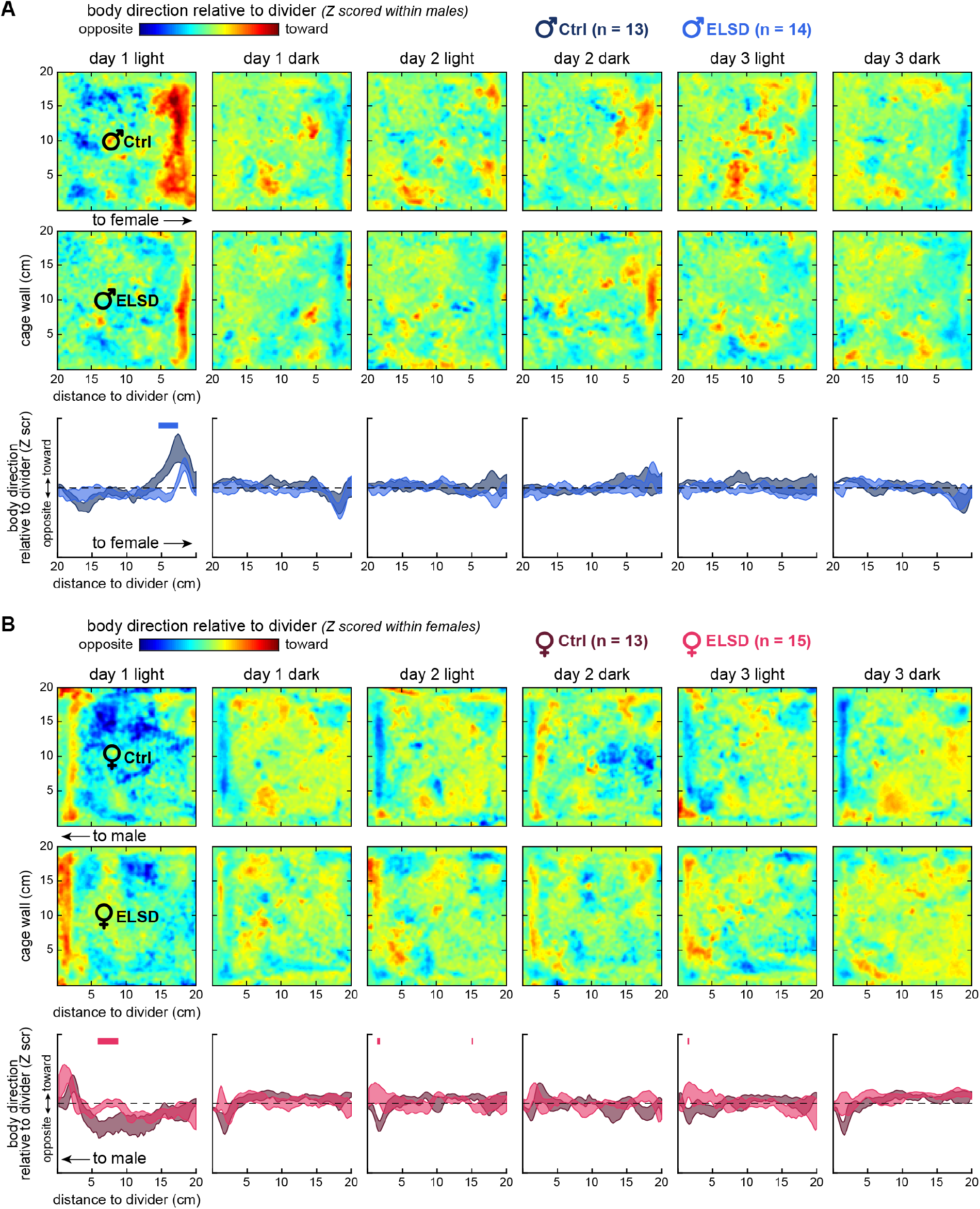
ELSD alters both spatial occupancy and body orientation in sex-specific manners. (A) Heatmaps: body direction was mapped onto home-cage quadrants (spatial axes, 2 mm grids) per recording period, and Z-scored across recording periods (see color code). Images are averages across the animals of each male group. Male-Ctrl animals spent more time near and toward the divider, especially during the light period of Day 1, as indicated by areas in red. Curves: heatmaps from individual animals and recording periods were averaged vertically, generating body direction vs. distance to divider curves. The standard errors in this panel represent variations from such curves. Male-ELSD animals were less likely to behave near and toward the divider on Day 1, light period. Blue bars on top indicate post-hoc differences at distance-to-divider bins (P < 0.05). (B) Same analysis, but from females. Female-ELSD animals were less likely to point opposite from the divider while occupying the center of the quadrant, particularly during the light period of Day 1. Across periods, Female-ELSD animals were slightly more likely than Female-Ctrl to behave near and toward the divider.

Figure 4B shows the same analysis, but from females. The stronger differences were again observed during the light period of Day 1. However, ELSD effects on female body direction were observed at a wider and more central location in the cage, 5-15 cm from the divider. Interestingly, at this location and day period, Female-Ctrl animals spent more time facing opposite from the divider than Female-ELSD animals (effect of ELSD vs. Ctrl grouping: F_1,2600_ = 52.734, P = 5.0*10^−13^) (Figure 4B; leftmost plots). No such effect had been observed in males (Figure 4A; leftmost plots). Thus, during the initial period of Day 1, here considered as the “novelty” period (Figures 2-3), higher behavioral activity enabled sex-specific body direction patterns across the home cage. These patterns could only be revealed due to the spatial mapping and Z scoring used in Figure 4, complementing Figures 2-3. We interpret these results as possibly sex-specific courtship displays, intermittent partner evaluation behaviors, or area preference patterns that are susceptible to ELSD. See also Movie S1.

Later recording periods in Figure 4B showed another difference between the female groups: Female-Ctrl animals were slightly more likely to orient themselves away from the divider when near it (Figure 4B; see narrow blue areas within 5 cm from the divider). This could reflect a behavior we sometimes observed during quiet wake or sleep periods, when animals rested their bodies against the divider, with their heads pointing slightly away from the divider, i.e., diagonally. We interpreted this behavior as possibly motivated by attempts to huddle with the other animal through the mesh divider, as this behavior was not regularly observed with any of the other 3 walls of the enclosure, which did not have a social stimulus on the other side. The difference between female groups in this putative huddling-like behavior was subtle and specific to the light period of Day 2 (group vs. distance interaction: F_99,2600_ = 1.311, P = 0.02), when Female-ELSD animals tended to rather point toward the divider when near it. These results shed light on both the sex specificity and ELSD sensitivity of subtle prairie vole behaviors.

## DISCUSSION

In this study, spatial and temporal analyses of prairie vole behavior over 72-hour cohabitation allowed us to make multiple new observations about pair bonding behavior with versus without ELSD. We both deepened previous ELSD male findings and found new effects of ELSD on female vole social behavior.

### ELSD affects social interactions primarily during the first 12 hours of cohabitation

We previously showed that ELSD during P14-P21 causes decreased huddling behavior during partner preference testing in adult male and female prairie voles and no preference to huddle with a pair bonded partner in males^16^. However, we had not previously described how ELSD affects social behavior during the cohabitation period, when the subjects are first exposed to a potential mate and the pair bond is formed. Here, we chose a cohabitation period of 72 hours across the mesh divider because preliminary data in our laboratory have shown this to be of sufficient duration for prairie voles to form a partner preference without mating during cohabitation. Furthermore, this cohabitation period is sufficient for males to induce estrus and sexual receptivity in female prairie voles^18,19^. Here, we describe how nuanced locomotor behavior relevant to socialization is altered in adult voles subjected to ELSD, compared to Control conditions: Male-ELSD animals showed signs of lower social interest in the initial 12 hours of cohabitation compared to control animals. This difference normalized over the subsequent 60 hours remaining during the cohabitation. It is still unknown whether this effect is due to the light phase in the first 12 hours, or if this effect would still remain if cohabitation was started in the dark phase. This could be examined in future studies. Regardless, the fact that Male-ELSD animals were significantly different from their controls is consistent with our previous findings that ELSD impairs social bond formation in male prairie voles^16^.

### ELSD affects behavioral rhythms across the 72-hour cohabitation

A serendipitous finding from our overhead tracking approach was that ELSD affected behavioral rhythms during cohabitation, compared to controls. Power spectral density analyses revealed that ELSD lowered the ultradian periodicity of male body direction independently on circadian rhythms, whereas in females ELSD disarranged these ultradian rhythms in a circadian-dependent manner. Ultradian and circadian activity patterns have recently been reported in prairie voles housed in a semi-natural enclosure: a 0.4 hectare field from which telemetry signals were used to examine periodical patterns of locomotor activity^20^. In addition, an older study described ultradian and circadian periodicity of wheel running in the common vole (*Microtus arvalis*)^21^. This indicates that the temporal organization of vole behavior – and probably of mammals in general – is robust enough to emerge in various experimental conditions, from the field to the laboratory. That was also the case in our home cage study, where behavioral activity was observed to be organized in cycles of 2-3 h. Our contribution, however, is to demonstrate that ultradian cycles are not only present in gross behavioral measures such as locomotion, but also in dissected behavioral components, such as body direction relative to the cage mate. Crucially, it is unlikely that the ultradian periodicity we found in body direction was merely a byproduct of general sex-unspecific behavioral rhythms, because ELSD effects were almost specific to the periodicity of body direction, in addition to being very different between males and females. This suggests that behavioral components may be orchestrated by separate “clocks”, each one involving sex-specific mechanisms with different susceptibilities to disturbances, such as ELSD. This notion is consistent with the common vole study mentioned above^21^, according to which wheel running, as well as feeding behavior, lose ultradian0020periodicity upon lesioning certain hypothalamic nuclei. Despite all of these findings, the purpose or predictive value of these fluctuations is unknown. Therefore, future vole studies involving manipulation of brain substrates relevant to social behaviors, such as the medial prefrontal cortex and its downstream circuits^22–24^, could be highly informative if combined in real time with behavioral tracking applications, like the ones described here, as well as physiological measures (e.g., cardiac activity and body temperature), which are also known to fluctuate in ultradian rhythms in voles^25,26^. Even simple correlational studies between such fluctuating behaviors and subsequent pair bonding may lead to novel insights or biomarkers for bonding.

### ELSD affects area occupancy patterns upon first meeting

Computer vision and our superimposed spatial analyses revealed a stereotyped locomotor pattern not previously described in prairie vole behavior involving body orientation and reorientation towards a social stimulus especially evident in the first 12 hours of cohabitation. While this behavior has not been previously reported in prairie voles, this physical orientation and reorientation towards the partner could be analogous to a stereotyped “social dance” of sorts – reported in other species, such as in Drosophila courtship behavior^27,28^. This behavior could also be the response to the mesh barrier interfering with typical male prairie vole courtship behavior^15,29,30^. Curiously, this “social dance” was significantly blunted in both ELSD males and females, compared to control animals. Notably this is also one of our findings in which ELSD females showed differences compared to controls. This speaks to the power and ability to detect nuanced behavioral differences when using an unbiased, high-resolution approach to behavioral tracking such as the custom analyses that we performed in these experiments. However, it should be noted that it is still unclear whether this locomotor pattern of activity would still be observed in response to a novel environment *without* a social stimulus. Further studies should examine the specificity of this activity pattern towards a social stimulus.

## Limitations

There are several limitations of this study, in addition to the caveats mentioned above. First, in our experimental design in which cohabitation of animals took place using a mesh divider, a major limitation is that this setup does not allow for assessment of true huddling, mating, or other fully interactive social behaviors. Now that our protocol has been successfully established, we could add further complexity by allowing more naturalistic interactions in future studies. Secondly, we did not test partner preference in a traditional way in these animals. We have previously shown in multiple prior studies a robust effect of ELSD on partner preference behavior, which is reasonably safe to generalize to these animals but should still be acknowledged as a caveat. Once we are able to allow further complexity and naturally interactive behaviors, it would be of great interest to apply our computer vision protocol to the partner preference test itself. Finally, it remains unclear whether the behaviors that we observe as most affected in ELSD animals are driven by being in the light phase, and/or by sleep, and/or by circadian processes. Dissecting these potential confounders is beyond the scope of the current study, but should be studied in the future.

## Conclusions

Early life sleep disruption has been shown to have long lasting effects on social behavior in adult prairie voles (*Microtus ochrogaster*), including impaired pair bonding with opposite sex mates during the conventional lab partner preference testing. Now, in this study, we used high resolution unbiased computer vision to identify and quantify novel, more nuanced social behaviors, including temporally distinct locomotor patterns of approach and orientation towards the partner, as well as ultradian social activity rhythms. ELSD males and females showed significant blunting of these behaviors compared to control animals in a temporally distinct manner across the 72-hour cohabitation period. Our findings highlight the utility of combining conventional manual behavior tracking together with unbiased computer vision approaches in order to paint a comprehensive picture of social behavior in animal models. Ultimately these more sophisticated approaches will aid in closing the gap in translation from preclinical to clinical studies, with the goal of understanding early life sleep effects on heterogeneous neurodevelopmental disorders such as autism.

## ACKNOWLEDGMENTS

We thank all members of Dr. Miranda M. Lim and Brendon O. Watson’s labs for internal discussions, especially rotation student Marcel Elkouri for his help during data collection. This work was supported by NSF NCS Foundations Award 1926818 and Portland VA Research Foundation to M.M.L., and R01MH126137 to B.O.W. The authors declare no competing interests.

## AUTHOR CONTRIBUTIONS

Conceptualization, B.O.W., C.E.J.T., L.S.B.J. and M.M.L.; Methodology, B.O.W., C.E.J.T., L.S.B.J., M.M.L. and P.T.W.; Software, L.S.B.J.; Formal Analysis, L.S.B.J.; Investigation, C.E.J.T., L.S.B.J. and P.T.W.L; Resources, B.O.W. and M.M.L.; Data Curation, C.E.J.T. and L.S.B.J.; Writing - Original Draft, L.S.B.J.; Writing - Review and Editing, B.O.W., C.E.J.T., L.S.B.J. and M.M.L.; Visualization, B.O.W., C.E.J.T., L.S.B.J. and M.M.L.; Supervision, B.O.W. and M.M.L.; Project administration, B.O.W. and M.M.L.; Funding Acquisition, B.O.W. and M.M.L.

## METHODS

### Subjects

Prairie voles were bred and reared by both parents at the Veterans Affairs Portland Health Care System. Litters with both males and females were submitted to early-life sleep disruption (ELSD) or Control conditions when pups were P14-P21 (Figure 1A; see below for ELSD procedure). Subjects were weaned at P21 into groups of 2-4 same-sex siblings per cage (Male-ELSD, Male-Ctrl, Female-ELSD, Female-Ctrl) and co-housed at the same breeding site until reaching adulthood. The groups of siblings were transferred between P50-P90 to the University of Michigan Medical School for the main recordings. Prairie voles were allowed to acclimate to the facility transfer for two weeks prior to experimentation. Housing conditions were the same throughout experiments, including controlled temperature, ventilation and humidity, bedding, ad libitum food (mixed diet of rabbit chow, corn, and cracked oats) and water (bottles/hydrogel), environmental enrichment (cotton nestlets and wooden blocks/sticks), and 14:10 h light/dark cycle (lights on at 5:00 am). Cages and nestlets were changed weekly. The prairie voles we used derived from a colony at Emory University (Dr. Larry Young), which in turn originated from field-caught animals in Illinois. Genetic diversity has been maintained through bi-annual donations among researchers across the USA (North Carolina State, University of California Davis, University of Colorado Boulder, and Florida State University). Procedures received bioethical approval from all institutions directly involved in this study (University of Michigan Medical School: PRO00009818; VA Portland Health Care System: 3652-18).

### Early-life sleep disruption

Litter-containing home cages with both parents were placed on an orbital shaker (turned on every 110 seconds for 10 seconds, 110 rotations per minute) when pups were at P14-P21 of age, thus generating ELSD phenotypes. Control animals were moved into the room with the shakers, but cages were not agitated (Figure 1A). Hydrogel was provided instead of water bottles during orbital shaking to prevent spillage in ELSD cages. Hydrogel was equivalently provided in Control cages. As described by our previous study^16^, ELSD is a gentle sleep disturbance method that predominantly affects infant REM sleep while preserving parental care and hormonal markers of stress.

### Cohabitation recording

Each adult individual emerging from the housing and sleep manipulation procedures described above was assigned to an opposite-sex individual. Pairs were formed randomly with all possible sex vs. sleep manipulation combinations: Male-ELSD/Female-ELSD, Male-ELSD/Female-Ctrl, Male-Ctrl/Female-ELSD, Male-Ctrl/Female-Ctrl. Male and female were then placed in bedded home cages (48.3 cm length, 25.4 cm width, 20.3 cm height), but separated from each other by a lab-made mesh divider, meaning that individuals could only roam within the confinements of their own quadrants (Figure 1B-C). Cage dividers were made with metal wire mesh (square mesh, 6.5 mm aperture, 22 cm wide, 28 cm high) covered on both sides with plastic sheet (clear polycarbonate, 0.5 mm thick) to prevent animals from climbing. The bottom rectangular portion of the mesh barrier was left exposed without the plastic sheet (6.5 cm high), allowing animals to exchange bedding and sniff each other. Crocs with chow/gel were placed uniformly across quadrants, with chow and gel being always positioned away from the divider, on the left and right sides of the animal, respectively. No environmental enrichment objects were placed in the quadrants. Finally, we placed cardboard barriers between neighboring cages, preventing male-female pairs from distracting each other. Animals were allowed to behave freely in their quadrants while being filmed from a 90° overhead angle for 72 h. For video recording, we customized a vibration-free optomechanical assembly (ThorLabs) holding two infrared-sensitive cameras (specifications below), each camera surrounded by four infrared illuminators (made in the lab from inexpensive LED boards). Two mesh-divided home cages were placed below each camera, allowing to record four male-female pairs at once (Figure 1B-C). This system was installed in a housing-approved room with circadian light/dark switching. Light/dark switching was innocuous to video brightness, as imaging was obtained with infrared reflectance.

We used two grayscale cameras (Basler, acA1300-60gm), each one attached to a fixed focal length lens (Edmund Optics, 6 mm UC Series). The cameras communicated via a network adapter (Intel Pro 1000/PT) with a host computer, and videos were written into an array of hard disks (RAID) for protected data storage. Cameras were configured in Pylon software (Basler) with 8-bit depth, 800-pixel frame width, 896-pixel frame height (no binning), 1328 kbps and 20-Hz frame rate. Exposure and brightness were adjusted by manipulating the lenses and infrared illuminators, without further adjustments to the camera software. Videos were acquired using StreamPix software (NorPix) into 6-h mp4 segments (H.264 codec) from the two cameras synchronously. Each camera recorded four quadrants, i.e., two male-female pairs. Quadrants were re-framed using Adobe Premiere and re-exported using Adobe Media Encoder, resulting in smaller videos with one individual per video (384-pixel frame width, 416-pixel frame height, 1025 kbps, 25-Hz frame rate, mp4 format, H.264 codec).

### Behavioral tracking

Re-framed videos were submitted to marker-less body tracking using DeepLabCut (DLC)^17^. We trained the DLC network to label seven body parts per individual: nose, left/right ears, shoulder, two locations along the back and tail base (Figure 1D). For network training, we used manually selected video frames representing a variety of scenarios: from clear imaging of the animal (no motion blur or obstruction of body parts) to challenging situations (e.g., with motion blur, curled posture when sleeping or eating, tail base hidden under bedding, nose hidden by the mesh divider when sniffing the cage mate). We then trained the network overnight using a lab server.

### Data analysis

Body part coordinates per video frame were saved as CSV files, imported to Matlab (Mathworks) and converted from pixels to centimeters. We then obtained three measures per video frame (Figure 1D) explained as follows. (1) Body direction relative to the divider: we fitted a line to the sequence of coordinates from tail to nose and determined the angle of the fitted line relative to the horizontal plane of the video frame. The angle was then rescaled from -1 (opposite from divider) to +1 (toward divider), with left and right directions treated equally. (2) Distance to divider: length between shoulder and divider on the horizontal axis of the video frame, regardless of the position on the vertical axis. (3) Locomotion speed: difference (hypotenuse) between the shoulder coordinates of frame n and frame n+1.

All measures were timestamped per video frame according to clock time in number of seconds x number of frames per second. For example, the first frame after 8:00 am on Day 1 of recording was identified as 720001 (8 h * 60 min * 60 s * 25 frames + 1 frame). Timestamping was made without restarting the clock at midnight, so that each frame could have a unique identifier across the 72-h recording. These timestamping procedures were used to align all recordings onto a common 72 h axis, given that all recordings were intentionally made with 15-30 min margins for later trimming. Clock time information was obtained from the file naming system of the video acquisition software (StreamPix, NorPix).

Body direction relative to divider, distance to divider and locomotion speed data per individual were averaged into 1 h bins (Figure 2A curves) or 20 min bins separated into the three recording days (Figure 2B curves). In either case, binned data were submitted to statistical comparisons between sexes or ELSD treatments per sex (two-way repeated measures ANOVA, followed by Tukey’s post-hoc comparisons at each time bin). The same data were averaged across 24-h periods (Figure 2B box plots) or light/dark periods of Day 1 (Figure 2C box plots) and submitted to one-way ANOVA.

The three behavioral measures were also examined for ultradian periodicity. Data were resampled from the original video frame rate to 1 s bins and analyzed using Welch’s power spectral density (PSD) estimate (6-h Hamming windows, frequency range of 0-3 cycles per hour in steps of 0.03 cycle). By examining the PSD curves, we found that peaks were mostly prominent within the 0.1-0.6 cycle/h band, which we interpreted to represent the ultradian fluctuations we observed in the raw data (Figure 3A-B curves). Thus, we summed PSD values within the 0.1-0.6 cycle/h band per individual (Figure 3B box plots) and did the same across behavioral variables and time periods (Figure 3C). Differences between groups and sexes were examined using two-way repeated measures ANOVA, followed by Tukey’s post-hoc comparisons at each period. We additionally subtracted ultradian power curves by their own moving averages with a sliding window of 3 data points (i.e., 3 periods) on a per-animal basis (Figure 3D curves). This resulted in curves with magnified light/dark alternation, representing circadian fluctuations. Data from light and dark periods were then separately averaged (Figure 3D box plots). Within-sex comparisons per light or dark period were made using one-way ANOVA.

Finally, using data in 1 s bins, we created spatial maps depicting both area occupancy and body direction (Figure 4). A 100 × 100 cell array was created in Matlab to represent a 2 mm grid of the home cage quadrant (quadrant dimensions: 20 × 20 cm). The cell array was cumulatively populated with body direction values across time bins, according to the animal’s position at each time bin. We then averaged the values per cell, which resulted in the maps. Six maps were created per individual, corresponding to the light/dark periods of Days 1-3. Such maps were arranged three-dimensionally, and Z scored across the 3rd dimension within males or females (represented by averages per group in the heatmaps of Figure 4). We then averaged each map vertically to obtain body direction vs. distance to divider curves. These curves were submitted to within-sex statistical comparisons (two-way repeated measures ANOVA, followed by Tukey’s post-hoc comparisons per spatial bin; see Figure 4 curves).

## REFERENCES

1. Buckley AW, Rodriguez AJ, Jennison K, et al. Rapid eye movement sleep percentage in children with autism compared with children with developmental delay and typical development. Arch Pediatr Adolesc Med. 2010;164(11). doi:10.1001/archpediatrics.2010.202

2. Simola P, Liukkonen K, Pitkäranta A, Pirinen T, Aronen ET. Psychosocial and somatic outcomes of sleep problems in children: a 4-year follow-up study: Outcomes of persistent sleep problems in children. Child Care Health Dev. 2014;40(1):60–67. doi:10.1111/j.1365-2214.2012.01412.x

3. MacDuffie KE, Shen MD, Dager SR, et al. Sleep onset problems and subcortical development in infants later diagnosed with autism spectrum disorder. Am J Psychiatry. 2020;177(6):518–525. doi:10.1176/appi.ajp.2019.19060666

4. Cohen S, Conduit R, Lockley SW, Rajaratnam SM, Cornish KM. The relationship between sleep and behavior in autism spectrum disorder (ASD): a review. J Neurodevelop Disord. 2014;6(1):44. doi:10.1186/1866-1955-6-44

5. McGraw LA, Young LJ. The prairie vole: an emerging model organism for understanding the social brain. Trends Neurosci. 2010;33(2):103–109. doi:10.1016/j.tins.2009.11.006

6. Kenkel WM, Gustison ML, Beery AK. A Neuroscientist’s Guide to the Vole. Curr Protoc. 2021;1(6). doi:10.1002/cpz1.175

7. Young KA, Gobrogge KL, Liu Y, Wang Z. The neurobiology of pair bonding: Insights from a socially monogamous rodent. Frontiers Neuroendocrinol. 2011;32(1):53–69. doi:10.1016/j.yfrne.2010.07.006

8. DeVries AC, Johnson CL, Carter CS. Familiarity and gender influence social preferences in prairie voles (Microtus ochrogaster). Can J Zool. 1997;75(2):295–301. doi:10.1139/z97-037

9. Williams JR, Catania KC, Carter CS. Development of partner preferences in female prairie voles (Microtus ochrogaster): The role of social and sexual experience. Horm Behav. 1992;26(3):339–349. doi:10.1016/0018-506X(92)90004-F

10. Insel TR, Preston S, Winslow JT. Mating in the monogamous male: Behavioral consequences. Physiol Behav. 1995;57(4):615–627. doi:10.1016/0031-9384(94)00362-9

11. Getz LL, Carter CS, Gavish L. The mating system of the prairie vole, Microtus ochrogaster: Field and laboratory evidence for pair-bonding. Behav Ecol Sociobiol. 1981;8(3):189–194. doi:10.1007/BF00299829

12. Beery AK, Christensen JD, Lee NS, Blandino KL. Specificity in sociality: Mice and prairie voles exhibit different patterns of peer affiliation. Front Behav Neurosci. 2018;12:50. doi:10.3389/fnbeh.2018.00050

13. DeVries AC, DeVries MB, Taymans SE, Carter CS. The effects of stress on social preferences are sexually dimorphic in prairie voles. Proc Natl Acad Sci U S A. 1996;93(21):11980–11984. doi:10.1073/pnas.93.21.11980

14. DeVries AC, DeVries MB, Taymans S, Carter CS. Modulation of pair bonding in female prairie voles (Microtus ochrogaster) by corticosterone. Proc Natl Acad Sci U S A. 1995;92(17):7744–7748. doi:10.1073/pnas.92.17.7744

15. DeVries AC, Carter CS. Sex differences in temporal parameters of partner preference in prairie voles (Microtus ochrogaster). Can J Zool. 1999;77(6):885–889. doi:10.1139/z99-054

16. Jones CE, Opel RA, Kaiser ME, et al. Early-life sleep disruption increases parvalbumin in primary somatosensory cortex and impairs social bonding in prairie voles. Sci Adv. 2019;5(1):eaav5188. doi:10.1126/sciadv.aav5188

17. Lauer J, Zhou M, Ye S, et al. Multi-animal pose estimation, identification and tracking with DeepLabCut. Nat Methods. 2022;19(4):496–504. doi:10.1038/s41592-022-01443-0

18. Witt DM, Carter CS, Carlstead K, Read LD. Sexual and social interactions preceding and during male-induced oestrus in prairie voles, Microtus ochrogaster. Anim Behav. 1988;36(5):1465–1471. doi:10.1016/S0003-3472(88)80217-3

19. Carter C, Witt DM, Schneider ZLH, Volkening D. Male stimuli are necessary for female sexual behavior and uterine growth in prairie voles (Microtus ochrogaster). Hormon Behav. 1987;21(1):74–82. doi:10.1016/0018-506X(87)90032-8

20. Wallace G, Elden M, Boucher R, Phelps S. An automated radiotelemetry system (ARTS) for monitoring small mammals. Methods Ecol Evol. 2022;13(5):976–986. doi:10.1111/2041-210X.13794

21. Gerkema MP, Groos GA, Daan S. Differential elimination of circadian and ultradian rhythmicity by hypothalamic lesions in the common vole, Microtus arvalis. J Biol Rhythms. 1990;5(2):81–95. doi:10.1177/074873049000500201

22. Jones CE, Chau AQ, Olson RJ, et al. Early life sleep disruption alters glutamate and dendritic spines in prefrontal cortex and impairs cognitive flexibility in prairie voles. Curr Res Neurobiol. 2021;2:100020. doi:10.1016/j.crneur.2021.100020

23. Amadei EA, Johnson ZV, Jun Kwon Y, et al. Dynamic corticostriatal activity biases social bonding in monogamous female prairie voles. Nature. 2017;546(7657):297–301. doi:10.1038/nature22381

24. Smeltzer MD, Curtis JT, Aragona BJ, Wang Z. Dopamine, oxytocin, and vasopressin receptor binding in the medial prefrontal cortex of monogamous and promiscuous voles. Neurosci Lett. 2006;394(2):146–151. doi:10.1016/j.neulet.2005.10.019

25. Lewis R, Curtis JT. Male prairie voles display cardiovascular dipping associated with an ultradian activity cycle. Physiol Behav. 2016;156:106–116. doi:10.1016/j.physbeh.2016.01.012

26. Beery AK, Loo TJ, Zucker I. Day length and estradiol affect same-sex affiliative behavior in the female meadow vole. Hormon Behav. 2008;54(1):153–159. doi:10.1016/j.yhbeh.2008.02.007

27. Spieth HT. Courtship behavior in Drosophila. Annu Rev Entomol. 1974;19(1):385–405. doi:10.1146/annurev.en.19.010174.002125

28. Kayser MS, Yue Z, Sehgal A. A critical period of sleep for development of courtship circuitry and behavior in Drosophila. Science. 2014;344(6181):269–274. doi:10.1126/science.1250553

29. Gavish L, Sue Carter C, Getz LL. Male-female interactions in prairie voles. Anim Behav. 1983;31(2):511–517. doi:10.1016/S0003-3472(83)80073-6

30. Graham BM, Solomon NG, Noe DA, Keane B. Male prairie voles with different avpr1a microsatellite lengths do not differ in courtship behaviour. Behav Processes. 2016;128:53–57. doi:10.1016/j.beproc.2016.04.006

